# Role of STING complex in differential retrograde signaling in cybrids with K versus H haplogroup mtDNA

**DOI:** 10.1101/402495

**Authors:** Kevin Schneider, Marilyn Chwa, Shari R. Atilano, Sonali Nashine, Nitin Udar, David S. Boyer, S. Michal Jazwinski, Michael V. Miceli, Anthony B. Nesburn, Baruch D. Kuppermann, M. Cristina Kenney

**Affiliations:** Department of Ophthalmology, Gavin Herbert Eye Institute, University of California Irvine, Irvine, CA 92697; Retina-Vitreous Associates Medical Group; Beverly Hills, CA 90211; Tulane Center for Aging and Department of Medicine, Tulane University, New Orleans, LA 70112; Cedars-Sinai Medical Center, Los Angeles, CA 90048; Department of Pathology and Laboratory Medicine, University of California Irvine, Irvine, CA 92697

## Abstract

Mitochondrial (mt) DNA haplogroups, defined by specific single nucleotide polymorphism (SNPs) patterns, represent populations of diverse geographic origins and may play a role in disparate disease susceptibilities found in different ethnic/racial populations. The most common European haplogroup is H, while the K haplogroup is highly associated with Ashkenazi Jewish populations. Studies using transmitochondrial cybrids (cell lines with identical nuclei but mitochondria from either H or K haplogroup subjects) demonstrated significant molecular and biological differences but mechanisms for these disparities are unclear. In this study, we hypothesized that there is differential retrograde signaling occurring between the Stimulator of Interferon Genes (STING) pathway and H versus K mtDNA haplogroups. Results showed that K cybrids exhibit increased levels of cytoplasmic mtDNA fragments. After STING Knock-Down, H cybrids had lower expression levels for *EGFR, BRCA1, DNMT3A*, *DNMT3B, HDAC1,* and *IFNα* genes, but upregulated *DNMT3A* compared to control H cybrids. The STING-KD K cybrids showed downregulation of *EGFR, DNMT3A, HDAC1, HCAD9, CFH*, and *CHI,* along with upregulation of *DNMT1* and *IL-6* compared to control K cybrids. Since all cybrids have identical nuclei, the STING DNA sensor system interacts differently with K haplogroup mtDNA compared to H mtDNA for genes related to cancer (*EGFR, BRCA1*), methylation (*DNMT1, DNMT3A, DNMT3B*), acetylation (*HDAC1, HDCA9*), complement (*CFH, CHI*) and inflammation (*IFNα, IL-6*). In summary, in non-pathologic conditions, (a) STING is an important retrograde signaling mechanism(s) and (b) cybrids possessing Ashkenazi Jewish mtDNA (K haplogroup) interact with the STING complex differently compared to H cybrids which affects various disease-related pathways.

## INTRODUCTION

Mitochondria (mt) possess unique circular DNA that is maternally inherited. The mtDNA encodes for 37 genes, including 13 protein subunits essential for oxidative phosphorylation (OXPHOS), 2 ribosomal RNAs and 22 transfer RNAs. (1-3) The non-coding region of 1121 nucleotides, known as the MT-Dloop, is critical for mtDNA replication and transcription. Recent studies report that small biologically active peptides called Humanin and MOTsC that encoded from the *16s* and *12s* rRNA regions of the mtDNA, respectively, are likely involved in various pathological processes. (4, 5) All cells have both nuclear and mitochondrial genomes contributing to disease processes. The transmitochondrial cybrids, which are cell lines with identical nuclei but the mtDNA from different subjects, have been used to identify the effects of an individual’s mtDNA upon cellular homeostasis.(6-9) Previous studies using transmitochondrial cybrids (cell lines with identical nuclei but mtDNA from either H or K haplogroup subjects) have shown that the K cybrids have (a) significantly lower mtDNA copy numbers, (b) higher expression levels for MT-DNA encoded genes critical for oxidative phosphorylation, (c) lower Spare Respiratory Capacity (SRC), (d) increased expression of inhibitors of the complement pathway and important inflammasome-related genes; (e) significantly higher levels of *APOE* transcription that were independent of methylation status; and (f) higher levels of resistance to amyloid-β_1-42_ peptides (active form) than the H haplogroup cybrids(10), but it has been unclear how the differential retrograde signaling occurs in H versus K cybrids. Previously, we have used the human retinal pigment epithelial (RPE) cybrid model to show that cybrids with K haplogroup mtDNA have (1) significantly increased expression of ApoE, a critical lipid transporter molecule associated with human diseases; (2) higher degree of protection from cytotoxic effects of amyloid-β_1-42_ (active form); (3) increased expression of inhibitors of the alternative complement pathways and important inflammation-related genes; and (4) elevated bioenergetic respiratory profiles compared to the H cybrids.(10) These findings suggest that an individual’s K haplogroup mtDNA contributes to lipid transport, cholesterol metabolism, complement activation and inflammation, factors critical for AMD, Alzheimer’s disease and other age-related diseases. However, the mechanisms of retrograde signaling by different mtDNA variants (K versus H haplogroup) to the nucleus are not known at this time.

It is recognized that diverse racial/ethnic populations have different risks for specific diseases. For example, African-Americans are susceptible to developing type 2 diabetes, obesity, prostate cancer and glaucoma. (11-14) and European-Caucasians are more prone to developing age-related macular degeneration (AMD), skin cancers, carotid artery disease and multiple sclerosis. (15, 16) The maternal origins of different human populations can be classified into haplogroups based upon the patterns of accumulated single nucleotide polymorphisms (SNPs) within the mtDNA. Either increased risk or protection for human diseases, including Alzheimer’s disease, AMD, cancers and diabetes, can be associated with the mtDNA haplogroup profile of the subjects. (2, 17-28)

The H haplogroups are the most common European mtDNA haplogroup, while the L haplogroups, representing individuals of maternal African-origin, are the oldest and most diversified haplogroup (www.MitoMap.com). The A12308G SNP defines the UK cluster that contains both the U and K haplogroups. The K haplogroups (also known as Uk) is further defined by the G9055A SNP, has a 1-6% worldwide distribution and represents approximately 10% of ancestral Europeans. Approximately 32% of Ashkenazi Jewish population is highly associated with the K haplogroup and can be classified into the K1a1b1a, K2a2a and K1a9 subsets. (15) The genetic profile of the Ashkenazi Jewish population has become more homogeneous because of limited numbers of founders, intermarriage within the group and population bottlenecks involving decreases in population sizes due to environmental and/or sociological events. (29)Behar, 2004 #7040} As a result, with respect to the genetic profiles, the Ashkenazi Jewish population is an excellent well-defined group for studies correlating genetics associations with specific diseases including hypercholesteremia, hyperlipidemia, cardiovascular disease, Gaucher disease type 1, Usher Type 3A, Tay-Sachs disease and *BRCA1/BRCA2* genes associated with breast and ovarian cancers. (30-34)

The STING (Stimulator of Interferon Genes) pathway represents a DNA sensor pathway used by cells to detect the presence of cytoplasmic DNA fragments, which then trigger activation of innate immune systems (Fig. 1). (35) The vast majority of research about STING has investigated the effects of viral and bacterial infections releasing intracellular, foreign DNA that activates the host defenses.(36, 37) STING can also be activated by “self-DNA” resulting in autoimmune disease such as Systemic Lupus Erythematosus (SLE) and Aicardi-Goutie syndrome.(38) It was recently shown that transfection of PCR-amplified mtDNA fragments into ARPE-19 cells induced inflammatory cytokines and the effects were blocked following knock-down of STING.(39) Interestingly, these effects were dependent on the size of the mtDNA fragments but not sequence or location. Previous studies have shown that different racial/ethnic mtDNA haplogroups are associated with varying disease susceptibilities and innate immune responses.(7, 10, 16, 40) To our knowledge, this is the first study to investigate whether the STING system is engaged in non-pathological signaling between the mtDNA and nuclear genome and whether using the cybrid model, the different haplogroups (e.g., H versus K) might elicit different responses in downstream genes from cells that have identical nuclei and culture conditions.

**Figure 1.**
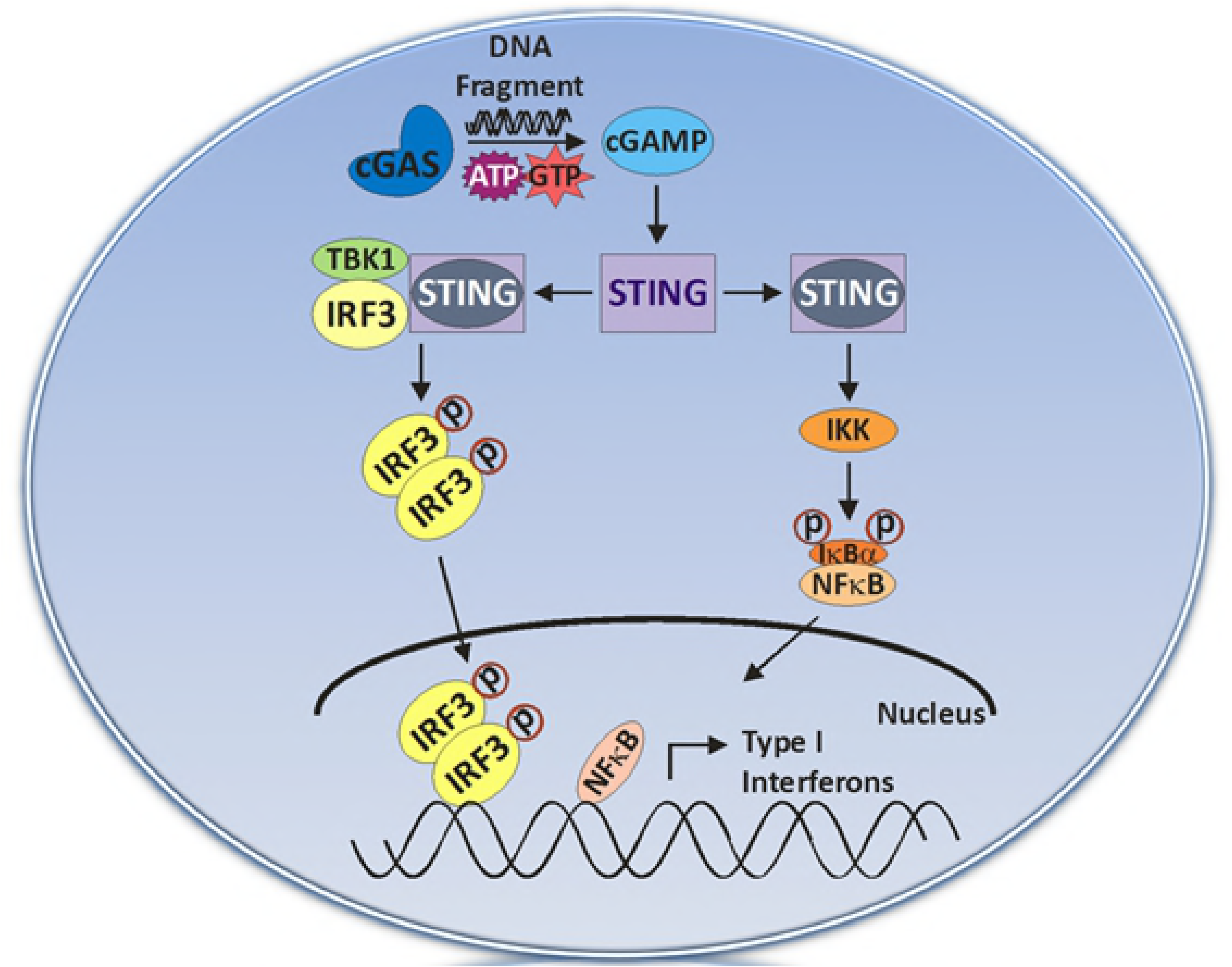
The STING (Stimulator of Interferon Genes) pathway.

## MATERIALS AND METHODS

### Generating Cybrid Cell Lines and Culture Conditions

Institutional review board approval was obtained from the University of California, Irvine (#2003-3131). There was no significant difference between the ages of the H subjects (n = 4, 42.5 ± 7.3 years) and K subjects (n = 5, 48.4 ± 3.6, p = 0.46) (Table 1).

**Table 1.** Subject information including cybrid numbers, gender, age, and haplogroups.

| **Cybrid** | **Gender** | **Age** | **Haplogroup** |
| --- | --- | --- | --- |
| **11-10** | **M** | **30** | **H4a1a** |
| **11-35** | **F** | **30** | **H1** |
| **13-52** | **F** | **58** | **H1** |
| **13-65** | **F** | **52** | **H41a1a** |
| **13-57** | **F** | **45** | **K1a1** |
| **13-65** | **F** | **38** | **K1a1b1a** |
| **13-75** | **F** | **56** | **K1c2** |
| **13-77** | **F** | **57** | **K1a1b1a** |
| **13-80** | **F** | **46** | **K1a1b2a1a** |

Peripheral blood was collected in sodium citrate tubes and DNA isolated using the DNA extraction kit (PUREGENE, Qiagen, Valencia, CA). Using a series of centrifugation steps, platelets were isolated, suspended in Tris buffer saline (TBS) and then fused with ARPE-19 cells that were deficient in mtDNA (Rho*0*) as described previously (9). Cybrids were cultured until confluent in DMEM-F12 containing 10% dialyzed fetal bovine serum, 100 unit/ml penicillin and 100 μg/ml streptomycin, 2.5 μg /ml fungizone, 50 μg/ml gentamycin and 17.5 mM glucose. All experiments used passage 5 cybrid cells.

### Protein extraction

H and K cybrid cell lines were plated in six-well plates for 48 hours. Cells were lysed using RIPA buffer (Cat. # 89900, Life Technologies), supernatants transferred to a new microfuge tube and concentrations of proteins were measured using Bio-Rad Dc protein assay system (Bio-Rad Laboratories, Richmond, CA, USA) according to the manufacturer’s instructions.

### Immunoblotting

Equal concentrations of total protein samples were loaded into the wells of 4–12% Bolt mini gels (Life Technologies) followed by SDS-PAGE electrophoresis. The gels were then transferred onto PVDF membranes. Following transfer, the membranes were blocked in 5% BSA/TBST for 1?hour, and incubated overnight at 4°C in primary antibodies. Blots were then washed with three times in TBST (Tris Buffered Saline-Tween20) and incubated with the respective secondary antibodies for 1?hour at room temperature. All primary and secondary antibodies were diluted in 5% BSA/TBST or 5% Milk/TBST as per manufacturer’s instructions. Following secondary antibody incubation, the blots were washed three times in TBST. Protein bands were detected using Clarity Western ECL Blotting Substrate (Cat. #1705060, Bio-Rad). β-actin antibody was used as a housekeeper protein control. Protein bands were visualized using Versadoc imager (Bio-Rad), and quantified using ImageJ software (NIH Image).

### Statistical Analyses

Data were subjected to statistical analysis by unpaired t-test, GraphPad Prism (Version 5.0, La Jolla, CA). P<0.05 was considered statistically significant. Error bars in the graphs represent SEM (standard error mean).

### Knock-down of STING

For siRNA mediated knockdown of STING, cybrid ARPE cells were seeded in 6-well plates at 7×10^5 cells/well. Thirty pmol final concentration of STING siRNA (#128591, ThermoFisher/Ambion,Waltham, MA) or Scramble siRNA were diluted in OPTI-MEM (ThermoFisher/Invitrogen) and incubated at room temperature for 5 minutes. Transfection reagent Lipofectamine 2000 (Invitrogen) was then mixed separately with OPTI-MEM as per manufacturer’s protocol and incubated for 5 minutes at room temperature. The OPTI-MEM/siRNA and OPTI-MEM/Lipofectamine tubes were then combined and incubation was carried out for 5 minutes at room temperature to allow formation of siRNA-lipid complex. Final mixture was then applied to cybrid cells in culture and allowed to incubate for 48 hours before RNA isolation.

### Isolation of RNA and Amplification of cDNA

RNA was isolated from untreated and STING-KD cultures (H cybrids, n=4; K cybrids, n=5) using the RNeasy Mini-Extraction kit (Qiagen) as described previously.(9) cDNA generated from 2 μg of individual RNA samples with the QuantiTect Reverse Transcription Kit (Qiagen) was used for qRT-PCR analyses.

### Quantitative Real-time PCR (qRT-PCR) Analyses

Total RNA was isolated from individual pellets of cultured haplogroup H cybrid cells (n=4 different individuals) and K cybrid cells (n=5 different individuals) as described above. qRT-PCR was performed on individual samples using QuantiFast SYBR Green PCR Kits (Qiagen) on an Applied Biosystems ViiA7 real time quantitative PCR detection system. Primers (QuantiTect Primer Assay, Qiagen or KicqStart Primers, Sigma) used to analyze for 37 different genes in various pathway: Complement *(CFH, CD59, CD55/DAF, CFI*); Methylation (*DNMT1, TRDMT1, DNMT3A, DNMT3B*); Acetylation (*HDAC1*, *HDAC2*, *HDAC3, HDAC4, HDAC6, HDAC9, HDAC10, HDAC11, HAT1*); Inflammation (*IL-6*, *IL-33*, *IL1β*, *IL-18*, *IFNα*, *IFNβ*; Chemokines (*CCL2*, *CCL20*); Cancer (*EGFR*, *BRCA1*, *ERBB2*, *ALK*, *PD1*); and STING pathway genes (*CGAS, TBK1, IRF3, IkBa, NFKB2, TRAF2, TNFRSF19*). Primers were standardized with the *HPRT1* or *HMBS* housekeeping genes. All analyses were performed in triplicate.

### Identification of Cytoplasmic DNA

Cells from cybrid cultures were collected (H cybrids, n = 4 and K cybrids, n = 5) and divided into two equal aliquots (1 × 10^6^ cells per aliquot). One set of aliquots was used for whole cell DNA extraction utilizing DNeasy Blood and Tissue kits (Qiagen) and these extracts served as normalization controls for the mtDNA copy numbers (see above). The second set of aliquots was resuspended in 500 μl buffer containing 50 mM HEPES (pH 7.4), 150 mM NaCl, and 25 μg/ml digitonin (MilliporeSigma, St. Louis, MO). The digitonin homogenates were incubated for ten minutes at room temperature on an end-over-end rocker to allow for selective plasma membrane permeabilization. Samples were then centrifuged at 1000*g* for three minutes to pellet intact cells. The supernatants were transferred to new tubes and spun at 17,000*g* for 10 minutes at room temperature to pellet any debris. This final spin yielded the cytosolic fraction. Cytoplasmic DNA was isolated and purified from this cytosolic fraction using the QIAQuick Nucleotide Removal Columns (Qiagen).

## RESULTS

### K haplogroup cybrids exhibit increased mitochondrial DNA fragments in the cytoplasm

Total cytoplasmic DNA was extracted from H and K cybrids (n=3) and analyzed for expression of both mitochondrial DNA markers (MT-ND2) and nuclear DNA markers (Actin). Cytoplasmic DNA content was then normalized to the total DNA. Mitochondrial DNA target expression was normalized to nuclear target expression, and overall cytoplasmic content normalized to total DNA levels (Fig. 2A). The K cybrids contained 4.3 fold higher levels of mitochondrial DNA in the cytoplasmic fraction compared to H cybrids. (p=0.049)

**Figure 2.**
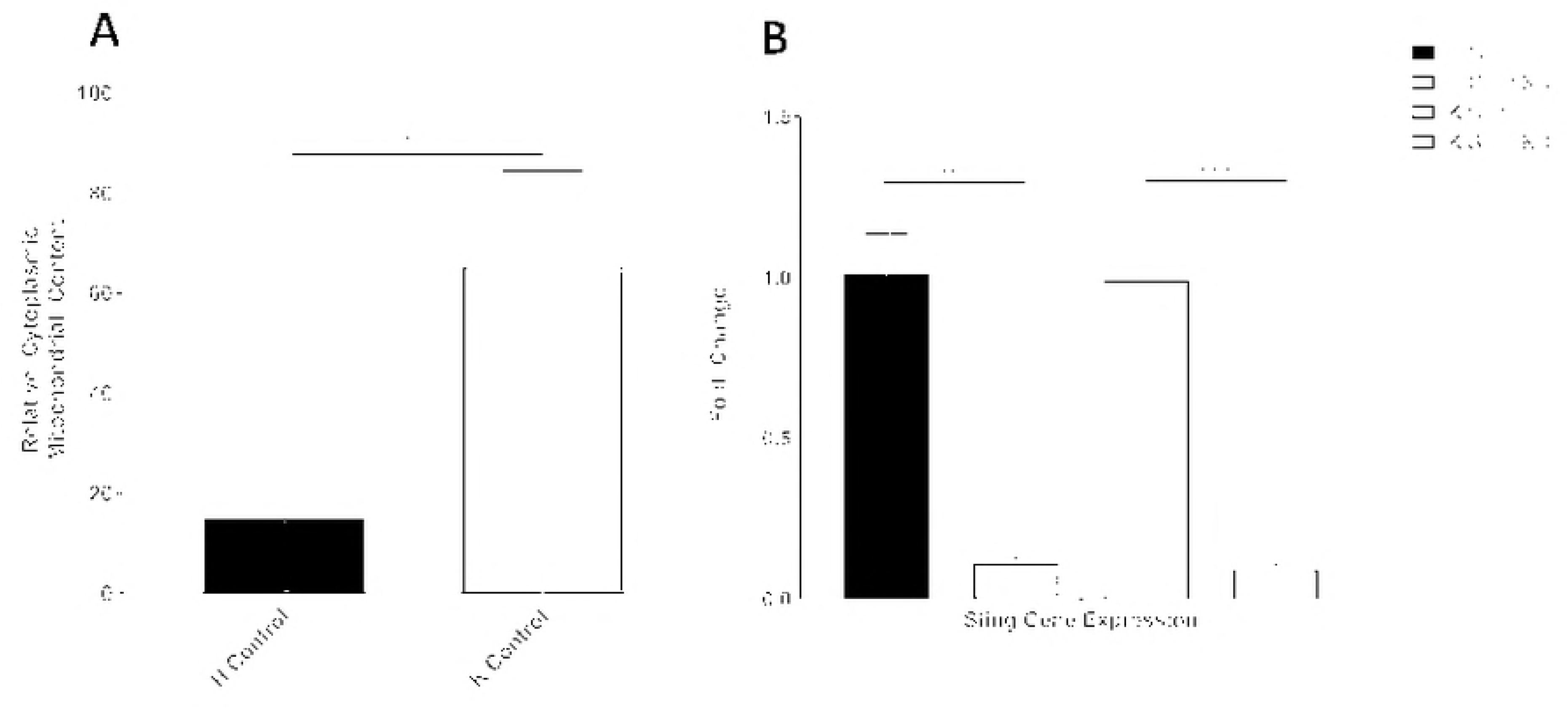
a) K haplogroup cybrids exhibit increased mitochondrial DNA fragments in the cytoplasm. b) Gene expression levels for STING at baseline were similar between H and K cybrids.

### Mitochondrial haplogroup does not affect STING gene expression

The STING complex is the intracellular sensor system for DNA fragments. Altered expression of downstream genes after STING knock-down is indicative that the DNA fragments are playing a role in the transcription for those genes. Gene expression levels for STING at baseline were similar between H and K cybrids (Fig. 2B). The H (n = 5) and K (n = 5) cybrids underwent STING knockdown (KD) by transfecting the cells with 30pmol siRNA or Silencer negative control. After 48 hours, RNA was isolated and the expression levels of the STING gene were measured by qRT-PCR. STING expression was significantly decreased in both the H and K cybrid cells. (11.48%, p=0.0028 in H cybrids and 9.5%, p<0.0001 in K cybrids; Fig. 2B). Gene expression was then analyzed for pathways related to cancer, epigenetics, complement and inflammation.

### Haplogroup K cybrids exhibit decreased expression of key DNA methylation genes

The K cybrids had lower levels of expression for DNMT1 (77.7% ± 4.3, p = 0.0057), *DNMT3B* (54.6% ± 3.4, p 0.0042) and *TRDMT1* (73.1% ± 4.2, p = 0.035), and DNMT3a (60.0% ± 2.7, p=0.0179) compared to the H cybrids. The levels for *MAT2B* were similar in the H and K cybrids (Fig. 3a). The expression levels for genes related to acetylation (*HDAC1, HDAC2*, *HDAC3*, *HDAC4*, *HDCA6*, *HDAC9*, and *HDCA11*) were similar in the H and K cybrids at baseline (Fig. 3b).

**Figure 3.**
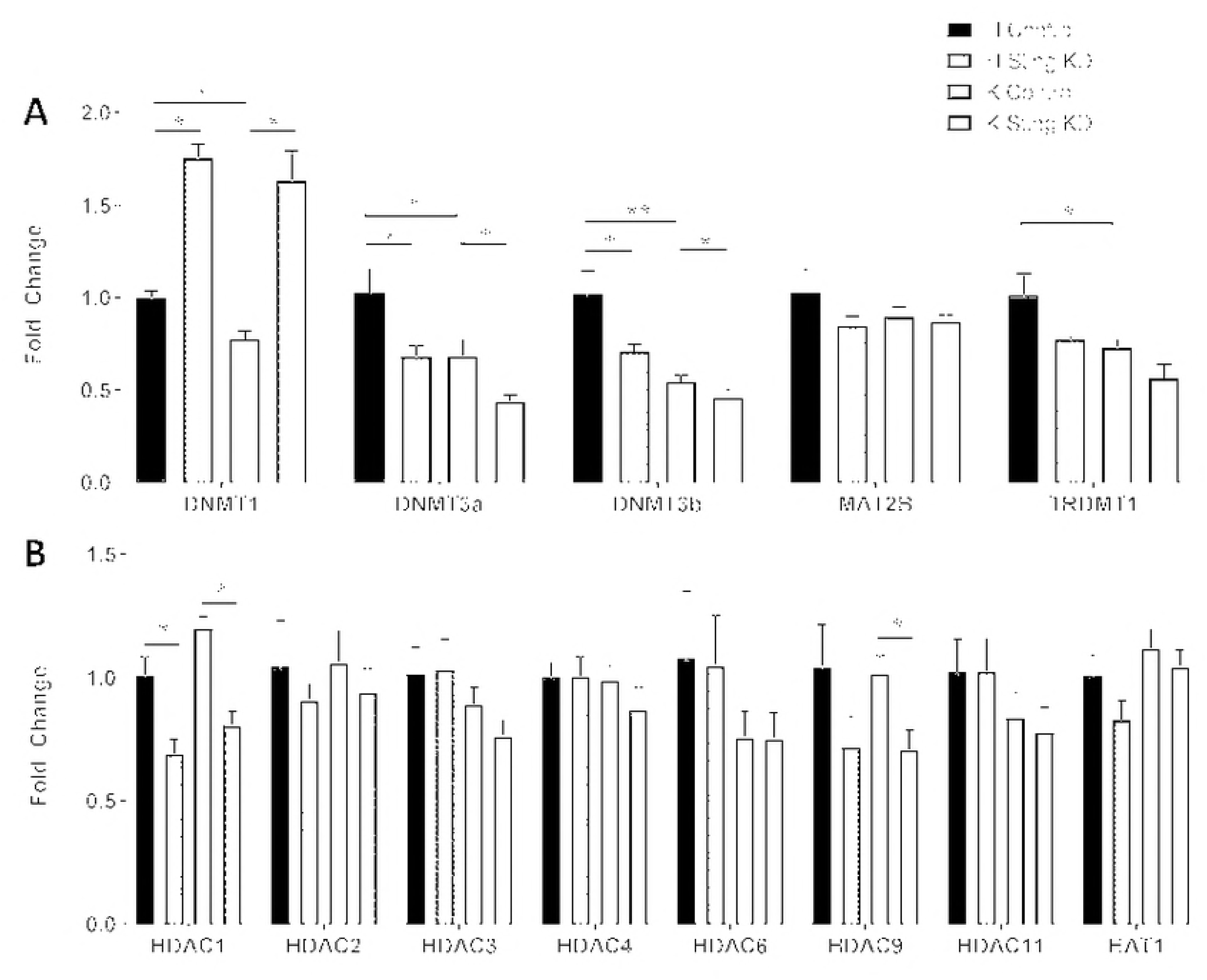
Haplogroup K cybrids exhibit decreased expression of key DNA methylation genes. a) The levels for *MAT2B* were similar in the H and K cybrids. b) The expression levels for genes related to acetylation (*HDAC1, HDAC2*, *HDAC3*, *HDAC4*, *HDCA6*, *HDAC9*, and *HDCA11*) were similar in the H and K cybrids at baseline.

### Knockdown of STING alters expression of epigenetic genes dependent on mitochondrial haplogroup

After STING-KD, *DNMT3A* gene expression dropped in both H and K cybrids (H: 68.2% ± 5.6, p=0.05; K: 43.9% ± 3.6, p=0.03) compared to the Control cybrids (Fig. 3a). *DNMT1* expression levels were increased significantly in both cybrids (H: 175.3% ± 7.4, p<0.0001; K: 163.4% ± 16.1, p=0.0009) compared to the Control H and K cybrids. For *DNMT3B*, the expression levels were lower in the STING-KD H cybrids (70.9% ± 3.9, p = 0.05) but not in the STING-KD K cybrids (p = 0.17). *TRDMT1 (DNMT2)* is a highly conserved methyl transferase that showed no change in expression after STING-KD in the H (p = 0.76) or K (p = 0.41) cybrids. The *HDAC1* expression decreased in both H and K cybrids after STING-KD (H: 69.2% ± 5.7, p=0.016; K: 80.8% ± 5.6, p=0.0009) versus Control cybrids (Fig.4b). The STING-KD K cybrid had lower levels of HDAC9 (70.9% ± 7.9, p = 0.02) while the expression levels in the STING-KD H cybrids were decreased compared to Control cybrids but was not significant (p = 0.18). STING-KD did not affect expression levels of *HDAC2*, *HDAC3*, *HDC4*, *HDAC6*, *HDAC9*, *HDAC10, HDAC11* and *HAT1* in the H or K cybrids (Fig. 3b).

### Haplogroup K cybrids differentially express markers of inflammation and RPE differentiation

The untreated K cybrids had lower levels of *IFNα* (54.2% ± 9.6, p = 0.006) and *CCL2* (24.6% ± 5.4, p = 0.0005) but higher levels of *IFNβ* (163.7% ± 12.2, p = 0.029) and *IL33* (185.3% ± 19.3, p = 0.04) compared to untreated H cybrids. The levels for *CCL20* (p = 0.08), *IL6* (p = 0.11), *IL1β* (p = 0.52) and *NLRP3* (p = 0.31) were similar in the untreated H and K cybrids.

### Knockdown of STING alters expression of inflammatory genes dependent on mitochondrial haplogroup

In the STING-KD K cybrids there was a decrease in *CFI* (72.6% ± 7.2, p=0.0045) and *CFH* (48.2% ± 3.1%, p=0.036; Fig. 4b). There was increased transcription levels in *IL6* (300.3.1% ± 44.6%, p=0.02) compared to K Control cybrids (160% ± 19.7%). The STING-KD H cybrids showed lower expression of *IFNα* (80.9% ± 5.3%, p = 0.04) compared to Control K cybrids. After STING-KD, there were no changes in expression levels for *CD55*, *CD59*, *CCL2*, *CCL20*, *IFNβ*, and *IL33* in either the H or K cybrids (Fig. 4a).

**Figure 4.**
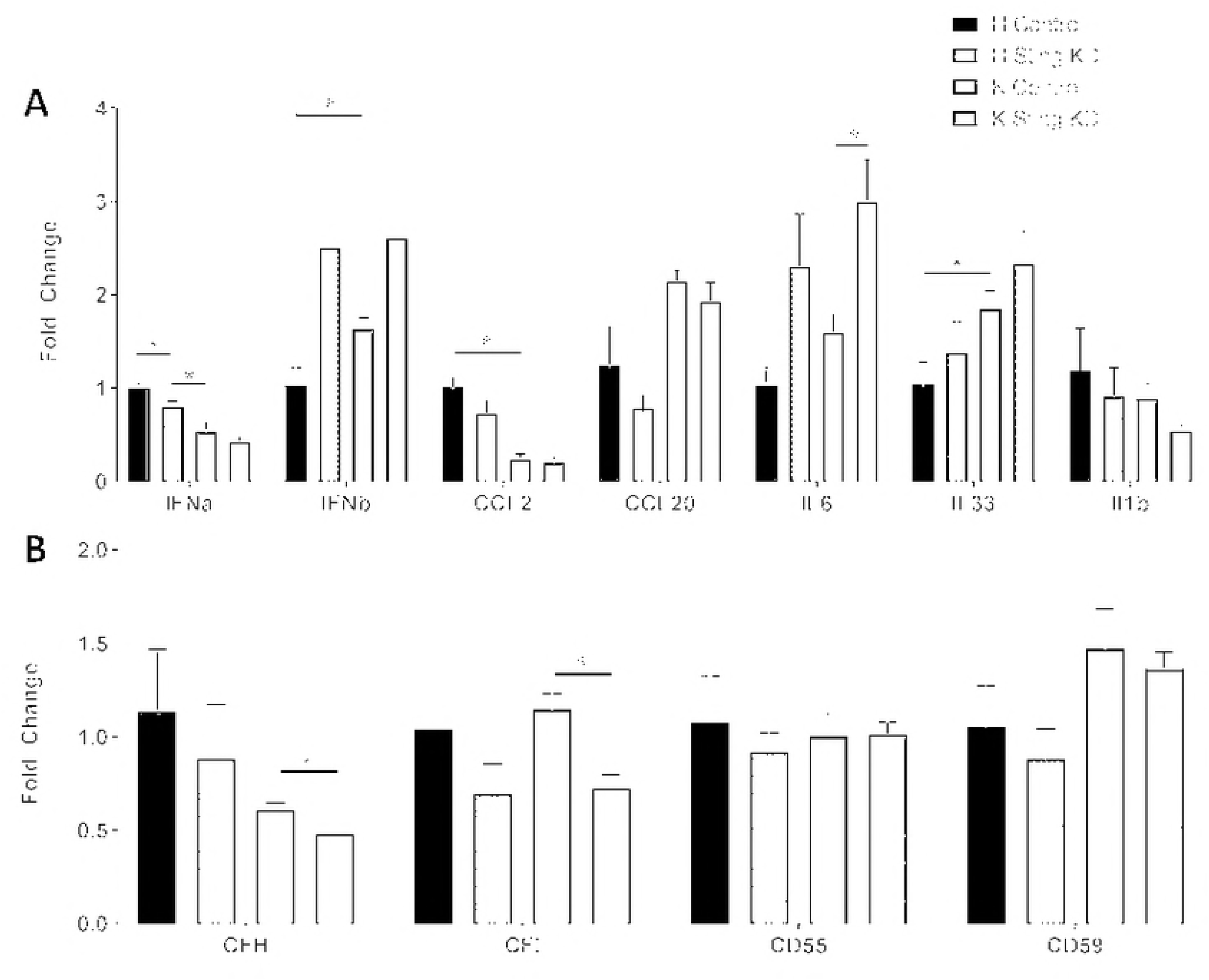
a) After STING-KD, there were no changes in expression levels for *CD55, CD59, CCL2, CCL20, IFNβ*, and IL33 in either the H or K cybrids. b) In the STING-KD K cybrids there was a decrease in *CFI* (72.6% ± 7.2, p=0.0045) and *CFH* (48.2% ± 3.1%, p=0.036).

### K haplogroup cybrids exhibit decreased expression of key cancer target genes

The cancer genes investigated in this study are known targets for drugs that are currently being used clinically to treat cancer patients (Table 2). The K cybrids had significantly lower expression levels of four cancer related genes (*BRCA1*, 51.6% ± 9.9%, p = 0.007; *EGFR*, 73.9% ± 6.9%, p = 0.05; *ALK*, 22.6% ± 5.9%, p = 0.003 and *PD1*, 40.5% ± 12.6%, p = 0.03) compare to H cybrids. The transcription levels for *ERBB2* (p = 0.3) were similar in H and K cybrids (Fig. 5a).

**Table 2.**
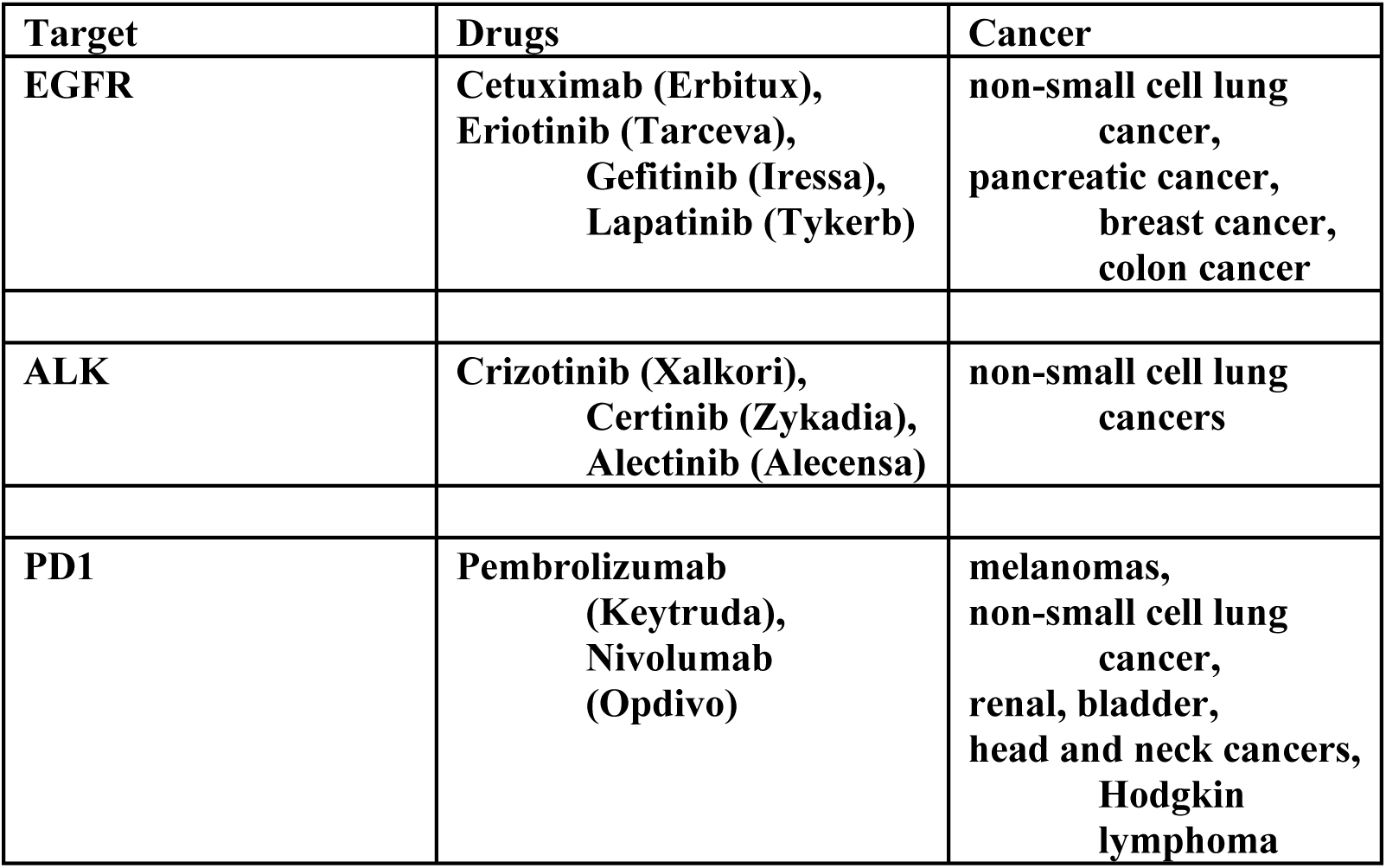
Description of Genes Targeted by Anti-cancer Drugs. The cancer genes investigated in this study are known targets for drugs that are currently being used clinically to treat cancer patients.

**Figure 5.**
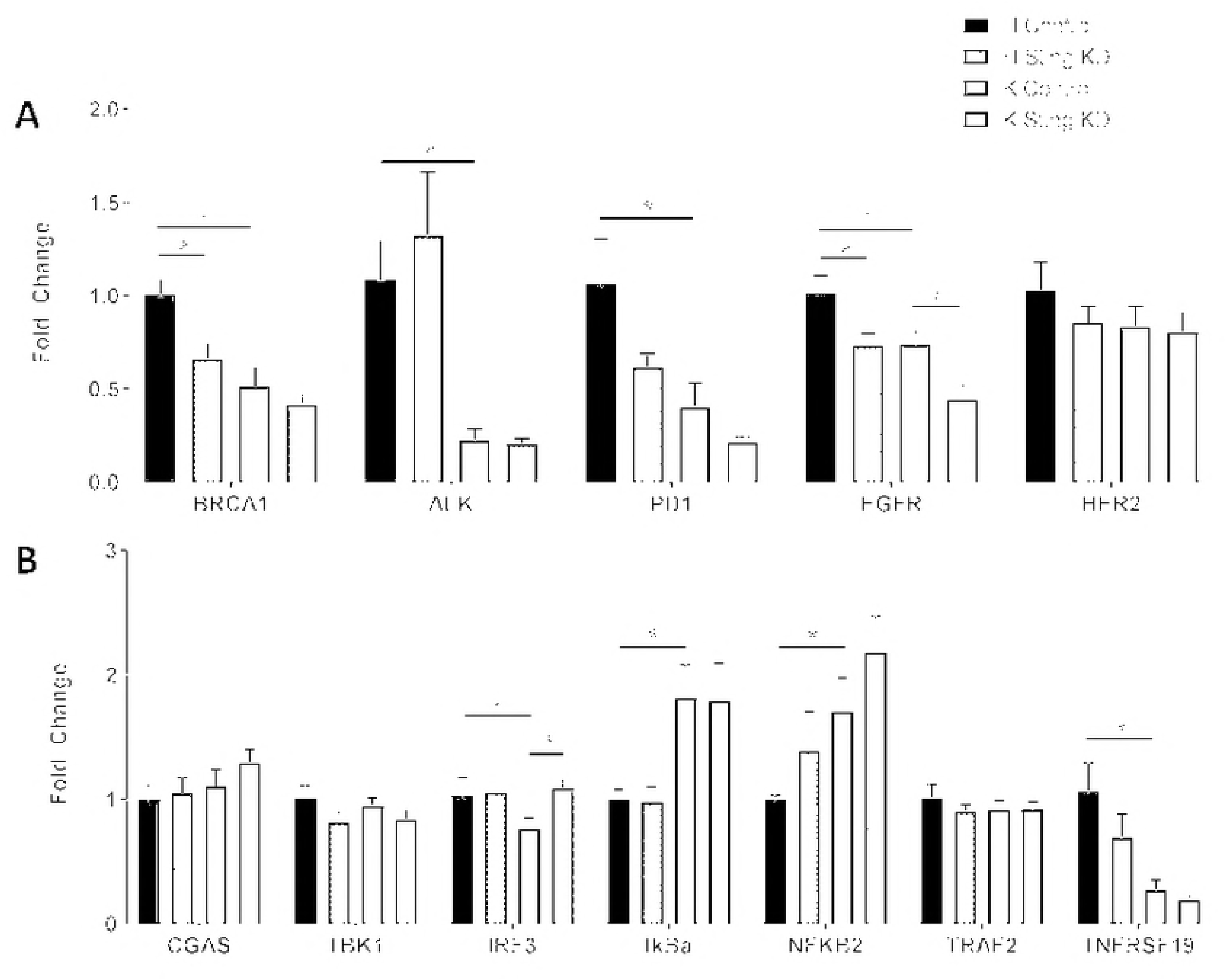
Gene expression levels in H and K cybrids. a) Knockdown of STING influences *BRCA1* and *EGFR* gene expression dependent on haplogroup and b) Knockdown of STING increases expression of *IRF3* in K haplogroup cybrids.

### Knockdown of STING influences BRCA1 and EGFR gene expression dependent on haplogroup

After STING-KD, the H cybrids showed lower *BRCA1* expression levels (34.7% ± 11.1%, p = 0.02) compared to H Control cybrids, while the K cybrid showed no significant decrease (p = 0.41). Conversely, the *EGFR* levels were lower in the STING-KD K cybrids (29.1% ± 10.1%, p = 0.02) and also in the STING-KD H cybrids (28.1% ± 11.8%, p = 0.05) compared to the untreated cybrids. The levels for *ALK*, *PD1*, and *ERBB2* were similar in the STING-KD versus Control H and K cybrids (Fig. 5a).

### K haplogroup cybrids differentially express genes involved in the STING DNA sensing pathway

A variety of genes involved in the STING signaling pathway were analyzed and we discovered that K cybrids had higher expression of *IkBa* (194.0% ± 30.2%, p = 0.05) and *NFKB2* (145.1% ± 11.9%, p = 0.026) as well as lower expression of *TNFRSF19* (27.5% ± 7.9%, p = 0.0073) and *IRF3* (67.9% ± 5.1%, p = 0.009) at baseline compared to H cybrids (Fig. 5b). At baseline there were no differences in expression of *CGAS*, *TBK1* or *TRAF2* between H and K cybrids.

### Knockdown of STING increases expression of *IRF3* in K haplogroup cybrids

After STING-KD, the only gene affected was *IRF3*, which was increased in the K cybrids compared to the lower baseline value (107.2% ± 8.7%, p = 0.0081). Interestingly, this increase returned *IRF3* gene expression to comparable levels of the untreated H cybrids. The levels of *CGAS, TBK1, TRAF2, IkBa, NFKB2*, and *TNFRSF19* were similar in STING-KD H and K cybrids (Fig. 5b).

### K haplogroup cybrids exhibit decreased phosphor-IRF protein levels

In order to confirm expression levels of key STING pathway genes and identify any alterations in STING signaling at baseline between the H and K cybrids, protein expression was measured via western blot. Phosphorylated and non-phosphorylated antibodies were used because many key STING pathway proteins function through phosphorylation. K cybrids demonstrated increased levels of IRF3 protein level (596% ± 58.3, p = 0.0015) and a decrease in the level of phospo-IRF3 (59.3% ± 10.8, p = 0.021) when compared to baseline H cybrids (Figs. 6a, 6b). No difference was seen in expression of NFKB or phosphor-NKFB (Figs. 6c-6e).

**Figure 6.**
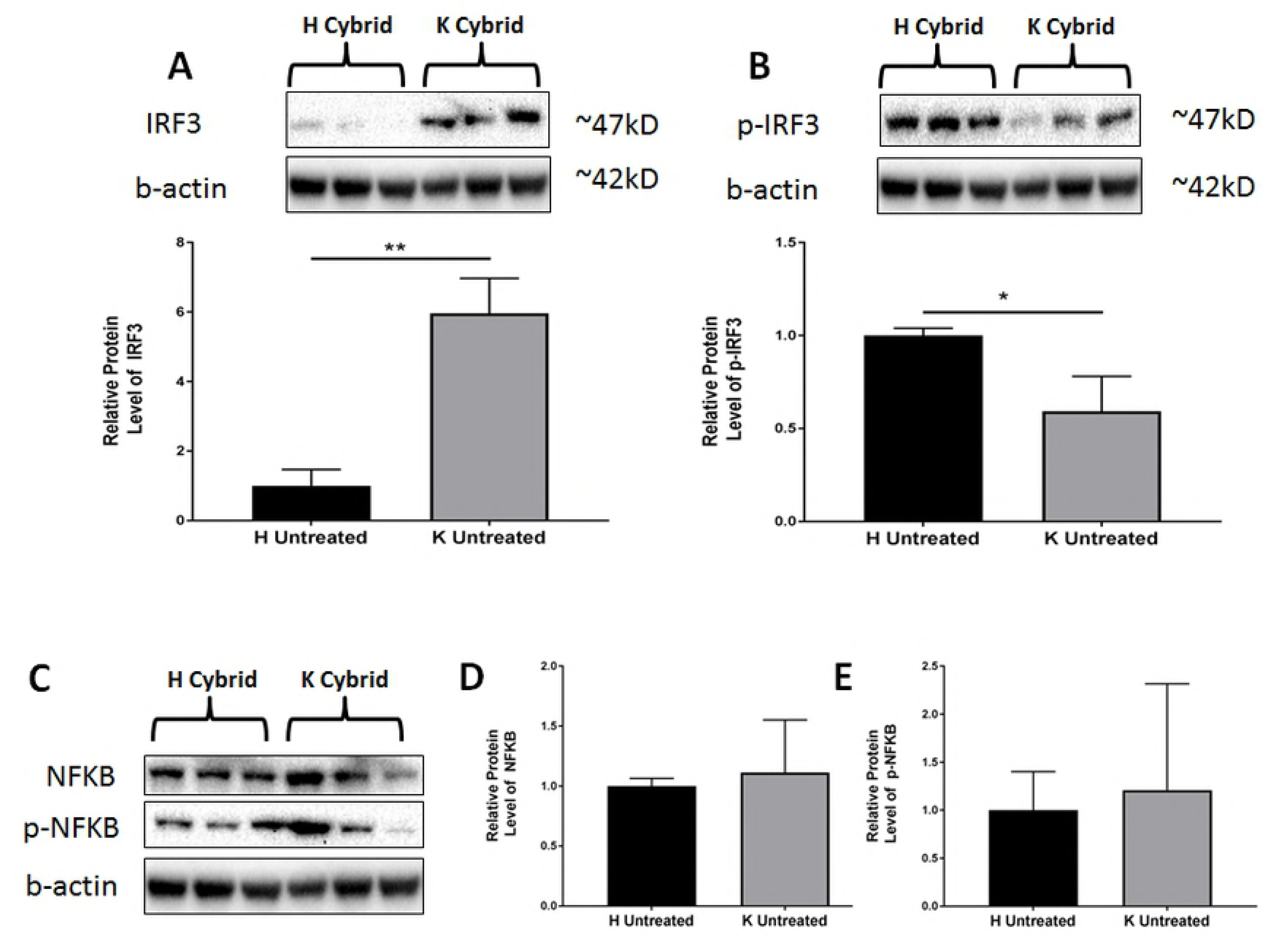
a, b) Phosphorylated and non-phosphorylated antibodies were used because many key STING pathway proteins function through phosphorylation. K cybrids demonstrated increased levels of IRF3 protein level (596% ± 58.3, p = 0.0015) and a decrease in the level of phospo-IRF3 (59.3% ± 10.8, p = 0.021) when compared to baseline H cybrids. c) No difference was seen in expression of NFKB or phosphor-NKFB.

## DISCUSSION

The present study was designed to determine if the STING (*TMEM-173*) pathway was involved in the signaling from the mitochondria to the nuclear genome in cybrids containing mitochondria from healthy subjects with either common European H haplogroup mtDNA or the Ashkenazi Jewish associated K haplogroup mtDNA. Studies have shown altered expression levels of genes associated with epigenetic pathways in K versus H cybrids and that inhibition of methylation with 5-aza-2’-deoxycytidine (5-aza-dC) altered the expression of NFκβ2, an important transcription factor activation of inflammation and immunity. (10)

The present study was designed to determine if activation of STING via DNA fragments played a role in the modulation of five methylation-related genes and eight acetylation-related genes. After STING-KD, both H and K cybrids showed increased expression of *DNMT1* and lower expression of *DNMT3A.* However, only the STING-KD H cybrids showed reduced *DNMT3B* expression levels while the STING-KD K cybrids were similar to control K cybrids. These findings indicate that the transcription of methylation pathway genes (*DNMT1, DNMT3A* and *DNMT3B*) can be regulated through STING, an intracellular DNA sensor system, and more importantly, the transcription levels are differentially expressed if the cells possess K haplogroup mtDNA compared to H haplogroup mtDNA. This potentially could lead to alterations in methylation patterns and variable modulation of downstream genes, depending if the subject has European mtDNA haplogroup versus Ashkenazi Jewish mtDNA profiles.

Of the eight acetylation genes investigated, only the *HDAC1* expression levels were reduced in both the H and K cybrids after STING-KD. This finding suggests that in non-pathological, unstressed cells there are some levels of STING activation via DNA fragmentation that upregulate HDAC1, a Class I histone deacetylase important for proliferation differentiation and apoptosis. This is the first description of a relationship between STING activation and modulation of HDACs. Interestingly, in the STING-KD K cybrids the *HDAC9* transcription was significantly lower (p = 0.02) while the STING-KD H cybrids showed a trend for lower *HDAC9* levels but it did not reach significant (p = 0.18) due to larger variations within the H cybrids. *HDAC9,* which is important for mitochondrial functions, has highest expression in brain (41) and has not previously been reported to be expressed in human RPE cells. *HDAC9* inhibits *Mef2* (myocyte enhancer factor2), which is important to oxidative phosphorylation in conventional T cells and T-regulatory (Treg) cells. (42) *HDAC9* also plays a role in Treg suppressive functions and inhibits transcription of *PGC1α* and *Sirt3*, both important for mitochondrial replication and ROS metabolism. These data suggest that via the STING pathway, the mtDNA can mediate the *HDAC9* expression levels, thereby influencing the mitochondria metabolism and possibly immune functions. However, additional studies are needed to more fully understand this relationship. The other six acetylation genes showed similar levels before and after STING knock-down.

The mechanism by which the STING complex affects the epigenetics is unknown. However, it has been shown that nuclear envelope transmembrane protein 23 (NET23)/STING can promote chromatin condensation and induce epigenetic changes, which is important because of its role in signaling for innate and apoptosis.(6) Green and coworkers reported that in response to viral-like double stranded RNA, the Pacific oyster (*Crassostrea gigas*) showed a upregulation of virus recognition receptors, signaling and effector genes, but the DNA methylation genes and STING remained unchanged, consistent with a poorly developed immune priming response. (43) This suggests that in mammalian cells, the activation of STING along with altered expression of specific methylation and acetylation genes may be important for immune recognition but the individual’s responses may be unique depending upon the ethnic/racial origin and underlying mtDNA profile.

It is recognized that pathological conditions (e.g., viral and bacterial infections) are often associated with DNA fragmentation and STING activation that modulates the immune responses. However, our findings suggest that STING activation may also be important for retrograde (mitochondria to nucleus) signaling under non-pathogenic conditions. In this study, the H and K haplogroup cybrid cell lines have identical nuclei and are cultured under non-stressed conditions. One can speculate that the fragmentation of the H mtDNA (European) versus the K mtDNA (Ashkenazi Jewish) might yield different size or variants of fragments that then activate the STING pathway differently, thereby leading to differences in the downstream regulation of the epigenetic genes. Our data shows that K haplogroup cybrids have increased levels of mtDNA in the cytoplasm of these cells. Due to the influence of this mtDNA load on the STING pathway, this could cause differences in methylation status, which might play a role in personalized responses to drugs and diseases that are often seen in the Ashkenazi Jewish populations. Using the cybrid model, higher levels of total global methylation have also been reported in cell lines with the European J haplogroup (44, 45), Ashkenazi Jewish K haplogroup (10), and also in cybrids with the African-origin L haplogroups (*unpublished data*) compared to those with the H haplogroup mtDNA. In turn, methylation patterns can influence the homeostasis of mitochondria, affecting apoptosis (46-48), and can be associated with disease susceptibilities and prognoses for cancers and age-related diseases. (49)

The field of therapies for cancer patient has been revolutionized by development of drugs targeting specific molecules key for progression and prognosis of the cancer. However, in spite of the use of these ‘targeting-drugs’ there are still many cancer victims that fail treatment and it is often not understood why they fail. In this study we analyzed the H versus K cybrids for the expression levels for 4 genes targeted by drugs; (a) Cetuximab, Erlotinib, Gefitinib and Lapatinib are inhibitors for *EGFR* production; (b) Crizotinibi, Ceritinib and Alectinib are inhibitors for *ALK*; (c) Pembrolizumab and Nivolumab are inhibitors of *PD1;* and (d) Pertuzumab and Trastuzumab targets the *ERBB2* (*HER2*) gene. We found that the untreated control H cybrids (European) had significantly higher expression of four of the genes (*EGFR*, p = 0.05; *BRCA1*, p = 0.007; *ALK*, p = 0.003; and *PD1*, p = 0.034) compared to the untreated K cybrids (Ashkenazi Jewish). After STING-KD for both H and K cybrids, the expression levels of *EGFR* were decreased (28.1%, p = 0.05 and 29.1%, p = 0.02, respectively) indicating that the *EGFR* is partially modulated via this DNA sensing system. Our findings suggest that the DNA fragments and STING pathway may be novel and previously unrecognized pathways to target for *EGFR* modulation in cancer patients.

If the cybrid findings are representative of what might be occurring in cancer patients, then there would be differential expression levels for these important genes, with European patients (H haplogroup) having higher transcription levels than Ashkenazi Jewish (K haplogroup) patients. Due to lower expression levels, K haplogroup patients might not respond equally to the inhibitor drugs for those gene products. This phenomenon may account for some of the differential responses found in clinical drug trials and also in prognosis outcomes for the cancer patients. It suggests that perhaps evaluation of the individual mtDNA profile may be of benefit to designing their treatment protocols. However, additional investigations are required fully understand the relationship between a person’s mtDNA haplogroup and their response to anti-cancer medications. In any case, this is the first report showing that the mtDNA variants can influence gene expression levels of these critical cancer-related genes.

The reported prevalence of BRCA1/2 mutation is has been reported to be higher in the Ashkenazi Jewish population (1 in 40) compared to other populations (1 in 400-800 persons). The expression levels of *BRCA1*, a gene highly associated with approximately 40% of inherited breast cancers and 80% of inherited breast and ovarian cancers, were also measured in K versus H cybrids. With respect to the *BRCA1* gene, the H cybrids had higher levels to begin with but showed a 34.7% decline after STING-KD (p = 0.02). In contrast, the K cybrids, with lower initial levels, were not affected by STING-KD. As *BRCA1* encodes for a tumor suppressor, individuals of the Ashkenazi Jewish population with the K haplogroup, may be expressing lower levels of this tumor inhibitor and therefore have less protection against cancer development.

Many of the studies related to the STING complexes are within the confines of viral infections. For example, it was reported that as herpes viruses induce mtDNA stress, the anti-viral signaling increases, leading to heightened responses of type I interferon. (36) While it is recognized that mitochondria are key participants in innate immunity required for robust anti-viral responses, our findings suggest that the STING sensor system is also a pathway of communication in healthy cells. This is not surprising as the symbiotic relationship between mitochondria and eukaryotic cells occurred over time and ancient signaling system would likely be maintained for cellular homeostasis (Fig. 6b).

K cybrids exhibit differential expression of several key genes associated with the STING pathway and its inflammatory response. Our data shows that K cybrids demonstrate increased gene expression of NFκβ2, an important gene associated with the inflammatory pathway of STING. This is consistent with the increased - mtDNA detected in the cytoplasm of K cybrids. However, western blot detected no difference of either NFκβ2 or phosphor-NFκβ2 levels between H and K cybrids (*data not shown*). Interestingly, there was decreased gene expression of *IRF3*, and decreased protein levels for phopho-*IRF3* in K cybrids. IRF3 and NκβB are critical to the STING pathway, which induces antiviral and pro-inflammatory cytokines, including type I interferons (IFN-α and IFN-β). Additionally, untreated K cybrids exhibited increased expression of *IkBa*, an inhibitor of NFκβ, but showed no difference in the expression of other key STING pathway related genes, such as *CGAS, TBK1* and *TRAF2*. The majority of data surrounding the STING complex has focused on its response to viral and bacterial infections, demonstrating that exogenous mtDNA can induce cellular inflammatory responses. However, our cybrid system is unique in that the mtDNA present in the cytoplasm is endogenous, rather than exogenous. Additionally, since the cells are under no external stress, the increased cytoplasmic content of mtDNA found in the K cybrids could be viewed as their normal state or non-pathogenic retrograde signaling.

The data from this study demonstrates that mitochondrial DNA haplogroup can exert a powerful effect on nuclear gene expression (Fig. 7a). One potential method by which mitochondrial DNA is able to elicit these changes is through the use of mitochondrial DNA fragments as signaling molecules (Fig. 7b). Most interestingly, there are subsets of genes that are differentially regulated by STING, dependent on mitochondrial haplogroup. This suggests that response to mtDNA fragments can be influenced by the mtDNA background. Additionally, this data demonstrates an association between mitochondrial haplogroup and mtDNA fragments levels. These fragments do not always illicit a pathogenic response, as would be expected. Finally, our data has identified novel pathways influenced by the expression of STING, suggesting that sensing of mtDNA fragments can have far reaching implications for signaling outside of inflammation.

**Figure 7.**
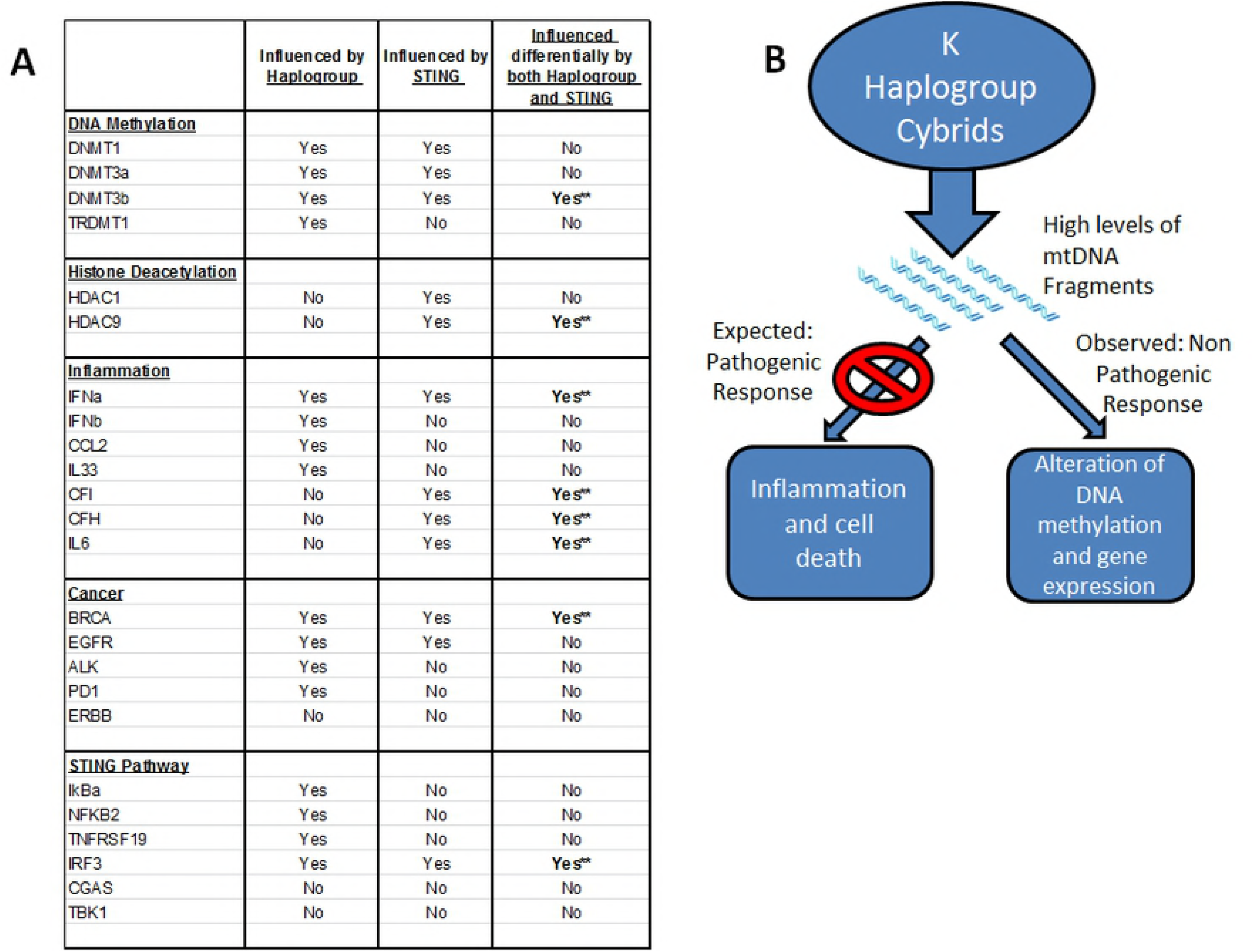
a) Nuclear gene expression influenced by haplogroup, STING and differentially by both Haplogroup and STING. b) One potential method by which mitochondrial DNA is able to elicit these changes is through the use of mitochondrial DNA fragments as signaling molecules..

## ACKNOWLEDGEMENTS

We wish to thank the subjects who participated in this study.

